# Whisker-based pre-neuronal and peripheral encoding of surface stickiness

**DOI:** 10.64898/2026.05.11.724292

**Authors:** Isis S. Wyche, Michaela A. O’Neil, Daniel H. O’Connor

## Abstract

Texture is a multidimensional perceptual feature of touch, with coarseness, stickiness, and compliance as its major axes of variability. Of these, coarseness is the best understood in the rodent whisker system. However, variation in surface stickiness is also a common feature of natural scenes, and is likely to alter the mechanical interactions between whiskers and surfaces that drive neuronal responses and are the basis for perceptual experience. In this study, we asked whether and how stickiness information could be extracted from whisker–surface interactions and represented in the activity of whisker follicle innervating mechanosensory neurons. We developed a 3D whisker tracking system applicable to texture sensing, and used it to characterize the whisker–surface interactions occurring during whisking against surfaces of jointly varying stickiness, coarseness, and position, as well as the responses of whisker follicle innervating neurons in the trigeminal ganglion. The bending, twisting, and roll of the whisker shaft, the rates and amplitudes of stick-slip events at the whisker tip, and the firing rates of a subset of mechanosensory neurons could all be used to distinguish between surfaces of high and low stickiness. These results demonstrate that stickiness information is available to the whisker system.

## Introduction

Texture is a ubiquitous aspect of tactile experience: any touch can provide information on the topography and material properties of the contacted surface. In the hand, this information supports grip control, dexterous handling of held objects, and object recognition in the absence of visual input. These are important behaviors in human daily life and for the study of sensory and motor function. The rodent whisker system is a well-characterized model for touch, including tactile texture discrimination. Mice and rats are capable of perceiving surface coarseness by whisker touch alone (Carvell & Simons, 1990; Isett et al., 2018; Morita et al., 2011; Zuo et al., 2011; Zuo & Diamond, 2019), distinguishing between gratings of varying spatial frequency or sandpapers of varying grade after a small number of exploratory touches using as little as a single whisker (Morita et al., 2011). This capacity may support whisker-touch-based exploration, navigation, and foraging behaviors. However, surface texture as perceived through the human fingertip is complex and multidimensional, including not only coarseness but also stickiness and compliance (Hollins et al., 2000; Okamoto et al., 2013; Picard et al., 2003).

These further dimensions of texture have been less studied in the whisker system, although they are associated with material properties that could affect the mechanics of whisker interactions with surfaces. It has not been determined whether any textural features beyond coarseness are perceived through whisker touch, or how they could be encoded. Clarifying these questions is important to the establishment of complete definitions of the perceptual space explored through whisker touch and the textural information represented throughout the whisker somatosensory system. We addressed surface stickiness as a step towards these goals.

An advantage of studying touch in the whisker system is that the whiskers can be tracked during contact with surfaces, allowing measurement of strains and estimation of stresses that are less accessible in hand and fingertip touch. The practical challenges of acquiring and analyzing video of whisker touch have generally limited whisker tracking to a single imaging plane (Pammer et al., 2013; Bush, Schroeder, et al., 2016; Campagner et al., 2016; Severson et al., 2017) or a pair of separately analyzed imaging planes (Hires et al., 2013). However, interactions between whiskers and objects are three-dimensional, and we anticipated that the complexity of whisker-surface interactions that could occur during sustained surface contact would necessitate three-dimensional consideration.

Changing curvature in whiskers pressed against an object (“bending”) is a prominent feature of whisker touch, and a significant driver of whisker follicle innervating neurons’ activity. While bending can be measured from the curvatures of both 2D and 3D tracked whiskers, 2D tracking captures only the apparent curvature within the imaging plane, potentially over- or under-estimating the true bending. The shape of the whisker may also deviate from planarity: the orientation in space of the plane in which it curves – its osculating plane – may vary with progression along the whisker, creating a helical shape in which both the curve’s curvature (the rate of its tangent vector’s rotation within the osculating plane) and its torsion (the rate of the osculating plane’s rotation about the tangent vector) are functions of arclength. Change in the torsion of the whisker shaft (“twisting”), which cannot be captured in 2D, is an additional strain that the whisker is subject to and that may drive neuronal responses. Finally, in addition to its changing shape, the whisker’s motion through space also has components not measurable in 2D, including transverse rotation and translation as well as roll about the longitudinal axis of the whisker follicle (Knutsen et al., 2008). Roll, as well, may be a source of strain in the whisker follicle able to drive responses in mechanosensory neurons.

Under some conditions, such as in the commonly used behavior of head-fixed whisking against a vertical pole (Bush, Schroeder, et al., 2016; Mehta et al., 2007; Campagner et al., 2016; O’Connor et al., 2010), the components of the whisker’s shape and motion that are not measurable in 2D can reasonably be treated as negligible. When significant torsion, roll, or out-of-imaging-plane curvature or motion of the whisker is present, however, 2D tracking cannot fully capture the stimuli capable of driving neuronal responses.

To maximize the explainable variability in neuronal responses to whisker touch requires a complete description of the three-dimensional stimuli they receive, which significant advances in the 3D tracking (Bush et al., 2021; Huet et al., 2015; Petersen et al., 2020) and modeling (Huet et al., 2022; Zweifel et al., 2021) of rodent whiskers have recently begun to provide. However, surface texture sensing has been outside the scope of existing tracking methods: it requires reconstruction of the full whisker as a non-planar curve during complex deformations, as well as full automation to accommodate the sub-millisecond resolution sampling needed to measure vibrations on the order of hundreds of hertz (Lottem & Azouz, 2009; Wolfe et al., 2008) and high-speed whisker micromotions of only milliseconds’ duration (Arabzadeh et al., 2005; Ritt et al., 2008; Wolfe et al., 2008; Oladazimi et al., 2021).

In this study, we aimed to extend our understanding of whisker-based texture sensing into the stickiness dimension. We developed a 3D whisker tracking method suitable for the investigation of texture sensing, and used it to characterize the effects of joint surface stickiness and coarseness variation on whisker–surface interactions and the responses of primary mechanosensory neurons innervating the whisker follicle. We found that whisking against surfaces of high stickiness was associated with broader sampling of the whisker bending, twisting, and roll space, stronger though less frequent stick-slip events at the whisker tip, and elevated spike rates in a subset of follicle-innervating neurons.

## Results

### Mice whisked against sequences of textured surfaces

To induce vigorous whisking, we trained water-restricted, head-fixed mice to run on a treadmill for water and sucrose rewards (Fig. 1A,B). Rewards were dispensed automatically when total running time since the previous reward reached 1.5-2.5 seconds, and sucrose solution was given instead of water on every third reward (Fig. 1B), to encourage voluntary running over extended periods under minimal water restriction. In each experiment that included electrophysiology, extracellular recording in the whisker region of the trigeminal ganglion (TG) captured the spiking of an isolated unit responsive to manual whisker deflection (Fig. 1C). Bouts of locomotion-induced whisking swept the whiskers across textured tiles placed within reach of each session’s whisker of interest, while a single high-speed camera captured simultaneous 4,000 fps video of the reflection of the whiskers and surfaces in top-down view silhouetted against an LED backlight (Fig. 1D) and the direct silhouettes of the whiskers and surfaces in rear view against a mirror reflecting the same backlight (Fig. 1E). The surfaces presented varied in material (Fig. 1F), spatial frequency (Fig. 1G), and position (Fig. 1H).

**Figure 1.**
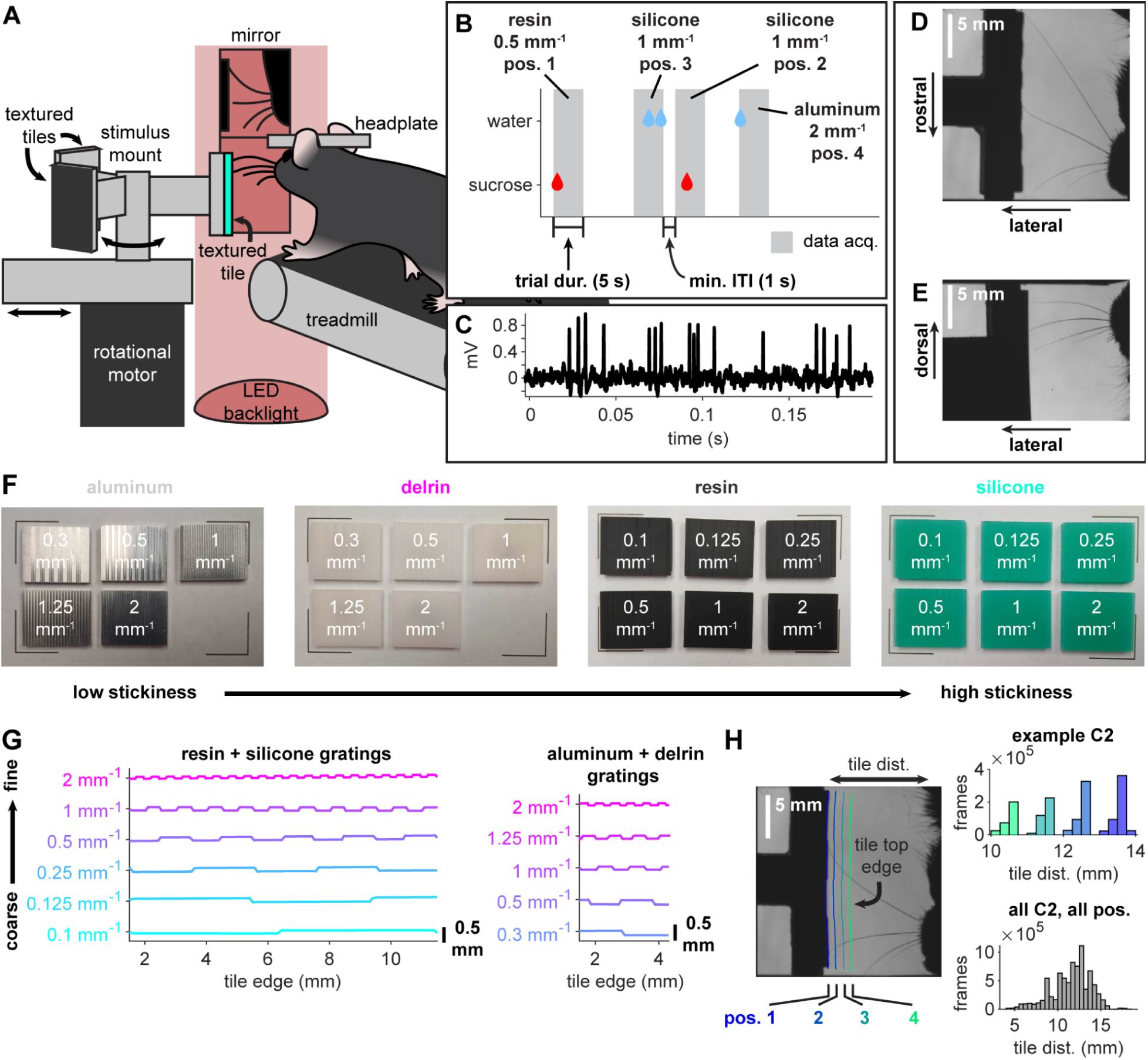
Mice whisked against surfaces of jointly varying stickiness, coarseness, and position, with simultaneous stereo videography and trigeminal ganglion electrophysiology. **A)** Experimental apparatus, viewed from the perspective of the camera. Some structural elements are excluded from the schematic for ease of visualization. B) Texture presentation sequence schematic. Vertical gray bars indicate trials; blue and red drops indicate times of water and sucrose solution dispensation, respectively. **C)** Unfiltered voltage trace showing spikes from an isolated whisker touch responsive unit in the trigeminal ganglion. D, E) Examples of top-down **(D)** and rear **(E)** views of whiskers and surface. **F)** Complete set of textured tiles used, grouped by material and ordered within material groups from coarse to fine spatial frequency (left to right, top to bottom). **G)** Grating waveforms used for 3D printer fabricated (resin and silicone, left) and CNC milled (aluminum and Delvin, right) tiles. **H)** Example tile positions in a single session (left and top right) and distributions of tile positions presented to all C2 whiskers in the dataset (bottom right).

In electrophysiological experiments, the whisker of interest was the single whisker in the isolated unit’s receptive field; otherwise, the identity of the whisker of interest progressed from C1 to C4 over successive experiments. All experiments included multiple whiskers, typically whiskers C1-C4; only one whisker in each experiment was the whisker of interest, but others could also make contact with the presented surfaces and were included in analyses. Whiskers grow rapidly prior to the quiescent phase of their growth cycle; to account for the possibility of growth from day to day when the same whisker appeared in multiple experiments, we treated each combination of observed whisker and observation date as a unique “whisker instance”.

Using the two-view video, we reconstructed the whiskers in 3D (Fig. 2A-C; Supplemental Videos 1-4) and extracted instantaneous whisker strain and kinematics from the reconstructions (Fig. 2D, E). In total, our dataset contains 88,560,000 samples (Fig. S1) of the shape and motion of 48 whisker instances, distributed over 23,100,000 video frames from 108 experimental conditions, 22 sessions, and 12 mice. Of these, 17,249,521 samples satisfied all criteria to be included in analyses: contact with the surface (closest tracked point ≤ 0.8 mm from the surface), active whisking (whisking amplitude ≥ 5°), and the absence of outliers in whisker strain or kinematics which we took to be indicative of tracking error.

**Figure 2.**
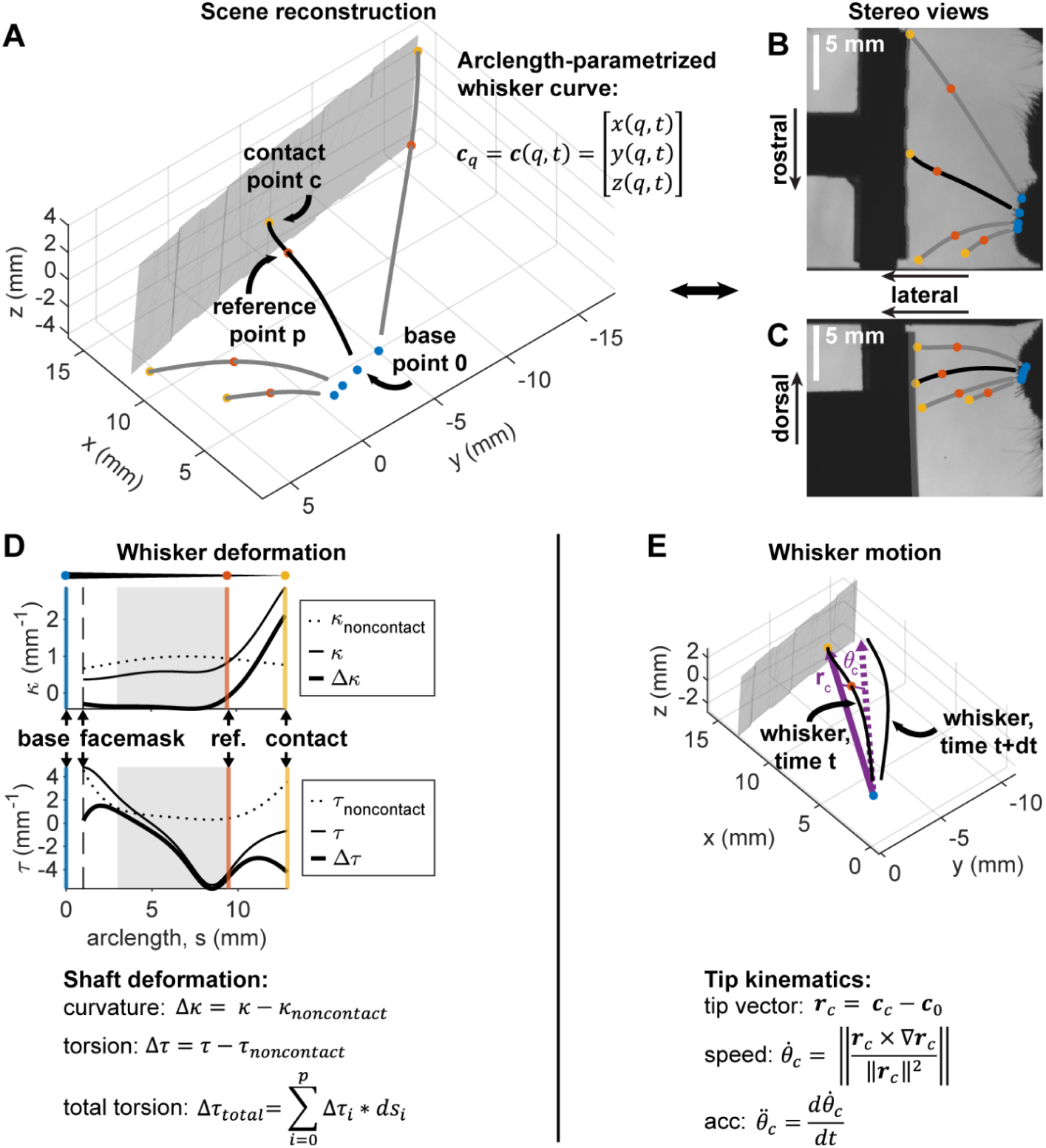
Whisker shape, strain, and kinematics were extracted from 3D reconstructions. **A)** 3D reconstruction of whiskers C1-C4 and the textured surface of a resin tile (gray mesh). Whisker centerlines (gray and black curves), estimated base positions (blue dots), mid-shaft reference points (orange dots), and tips or whisker–surface contact points (yellow dots) are shown. Whisker centerlines end at their intersections with the face silhouette masks (not displayed). **B, C)** Reprojection of whiskers and points in A into original top-down (B) and rear (C) views. **D)** Curvature and torsion evaluated along the full whisker shaft for the example frame in A. The measured curvature and torsion and the estimated noncontact curvature and torsion given the whisker’s position in this frame are used to compute the curvature and torsion deformations at each point. Gray shading indicates the region over which the change in torsion Δτ is integrated to obtain the total twisting Δτ_total_; vertical lines indicate the arclengths of the estimated whisker base (zero), the whisker’s intersection with the facemask, the fixed mid-shaft reference point, and the contact point. **E)** Computation of whisker tip kinematics. Whisker C2 is displayed at times t and t + dt; for visualization, dt is 80 ms rather than the true frame period of 0.25 ms. The angle between the base-to-tip vector r_c_(t) (solid purple vector) and r_c_(t+dt) (dashed purple vector) is θ_c_.

### Three-dimensional whisker tracking to capture stimuli not measurable in 2D

To quantify a whisker’s instantaneous shape, we computed its centerline’s total curvature and torsion in 3D space. As 2D analogs to the 3D curvature, we also computed the total curvature in the top-down imaging plane and the total curvature in a plane orthogonal to the top-down imaging plane. Torsion has no 2D analog, as a description of a plane’s rotation in space, but we considered 2D curvature a loose comparator for torsion as well.

When at rest and not in contact with a surface, the whiskers were nearly planar, with near-zero torsion over most of the shaft (Fig. S2A). However, torsion increased as the whisker twisted out of its intrinsic shape during contact (Fig. S2B). During whisking against surfaces, torsion and all three curvatures varied periodically, with amplitudes having the same order of magnitude (Fig. S2C,D). The amplitude of curvature variation in the top-down imaging plane did not overwhelm those of torsion in 3D space or curvature out of the top-down imaging plane. This suggests that the components of whisker shape not measurable in a top-down view should not be considered negligible on the basis of amplitude.

As measures of whisker motion, we computed the whisker base vector’s transverse rotational speeds in 3D, in the top-down imaging plane, and in the plane orthogonal to the top-down imaging plane, as well as the speed of the 3D whisker’s roll out of the top-down imaging plane. Like torsion, roll has no 2D equivalent, but we considered the speed of the whisker base’s rotation in the top-down imaging plane a loose comparator for roll speed.

As with the whisker shape variables, roll speed and all three transverse rotational speeds varied periodically during whisking against surfaces (Fig. S2E). The three transverse rotational speeds varied with amplitudes having the same order of magnitude, while roll speed amplitude was only one order of magnitude lower, indicating that the components of whisker motion not measurable in a top-down view should also not be considered negligible based on their amplitudes (Fig. S2F).

A second basis for considering a signal to be negligible is redundancy, which we quantified as the coefficient of uncertainty between a shape or kinematic variable measured in 3D and its 2D comparator in the top-down imaging plane. The coefficient of uncertainty between two random variables is a normalized mutual information value, representing the fraction by which knowing the value of one variable reduces uncertainty about the value of the other. By this measure, 3D whisker curvature and torsion were not redundant to curvature in the top-down imaging plane: curvature in the top-down imaging plane generally reduced uncertainty about 3D curvature and torsion by less than half (Fig. S2G). Whisker roll speed and 3D transverse rotational speed were likewise non-redundant to transverse rotational speed in the top-down view.

Finally, we noted that the motion of the whisker tip was not well constrained to the top-down imaging plane (Fig. S2H). The whisker tip traversed complex trajectories over the surface, including motion out of the top-down imaging plane that, if only that view were available, could appear as a slowing or stopping of whisker motion. Stick-slip events in the whisker’s motion over a surface are detected as characteristic tip speed, acceleration, and jerk transients (Ritt et al., 2008; Wolfe et al., 2008; Oladazimi et al., 2021); if these signals do not capture the true kinematics of the whisker tip, stick-slips may be missed, or periods of continuous high-speed motion may be incorrectly detected as containing stick-slips.

These observations demonstrate that during whisking against surfaces, 2D tracking may not produce reliable measurements of whisker strain or motion, and may miss or incorrectly detect stick-slips at the whisker tip. When this is the case, estimates of the relationships between neuronal activity and whisker strain, whisker stress, or stick-slip events also become less reliable.

### Whisking against the stickiest material increased sampling of high-strain whisker states

During whisking against surfaces, the whisker shaft deforms under the combined influences of the whisker tip’s interaction with the surface and the driving motion of whisking applied to the base. By altering the coefficient of friction between whisker and surface, changing the boundary condition at that interface, we expected that varying the material whisked against would also alter the whisker deformations produced.

We defined contact-induced whisker deformation as the whisker shaft’s bending and twisting out of its intrinsic shape and roll out of its intrinsic orientation. Whisk cycles typically traversed the contact-induced bending, twisting, and roll space as C-shaped trajectories (Fig. 3A). To quantify the effects of surface material on whisker deformation, we reduced the whisker deformation time series measured for each whisker instance to probability distributions conditioned on the material, spatial frequency, and position of the surface whisked against (Fig. 3B). A change in the sampling of the whisker deformation space between experimental conditions could appear as a change in the dispersion of the distribution, measured as the volume of its 50% highest density region (Fig. S3A); a change in its central tendency, measured as its expected value; or a combination of both effects.

**Figure 3.**
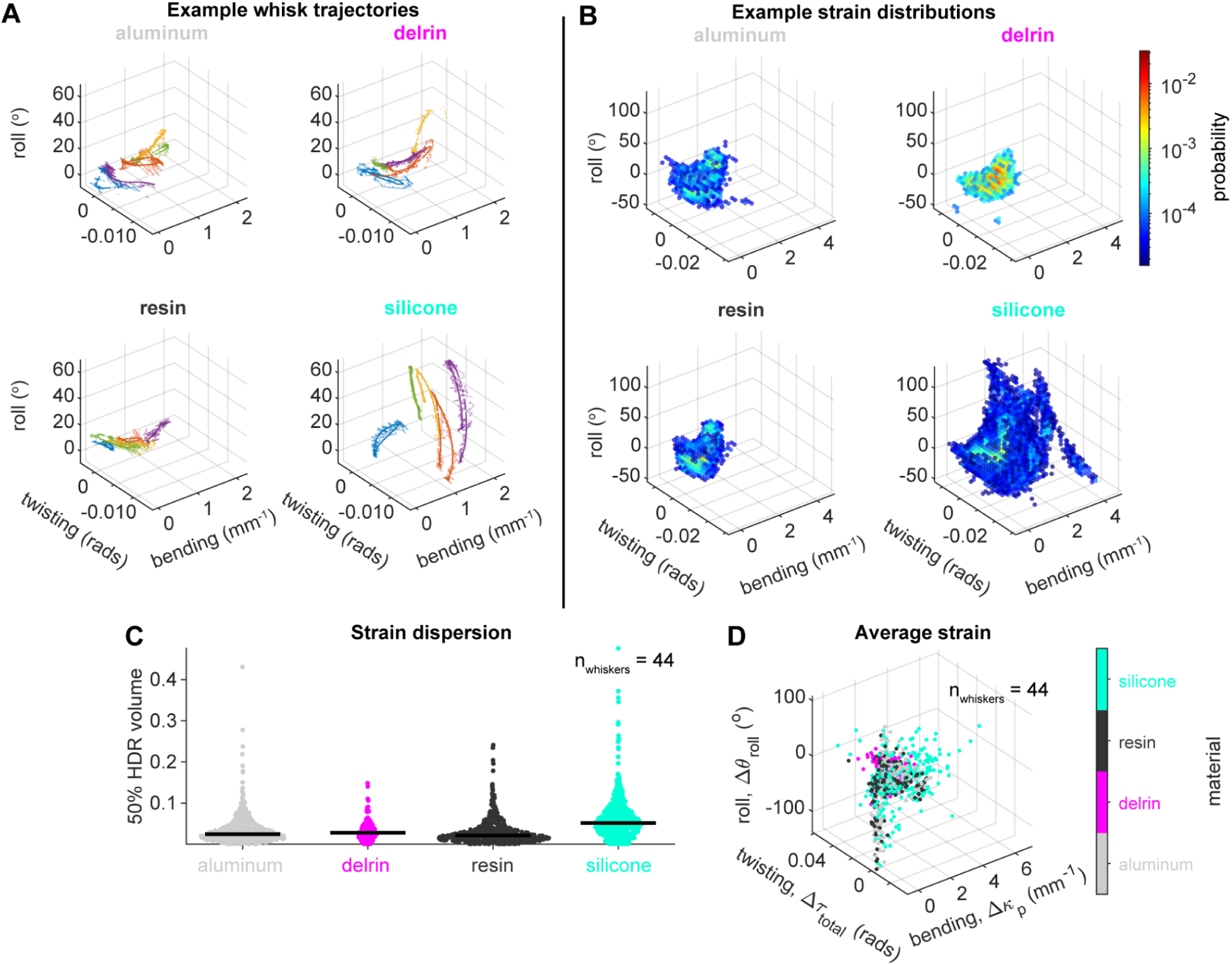
Whisking against silicone sampled a broader region of the whisker strain space than whisking against less sticky surfaces, reaching greater magnitudes of strain. **A)** Trajectories through bending, twisting, and roll space of five example whisks (colors) against each material. Opaque lines are smoothed data; transparent lines are unsmoothed. **B)** Example whisker deformation distributions for whisking against each material. **C)** Scatter plots of the 50% highest density region volumes of all whisker deformation distributions, grouped by material. Each point corresponds to a single whisker instance observed under a single experimental condition (a unique combination of surface material, grating frequency, and position). **D)** Mean whisker deformation values for all whisker instances in C, color coded by material.

Across all surface positions and spatial frequencies, we observed an increase in the dispersions of the whisker deformation distributions measured during whisking against silicone compared to the other, less sticky materials (aluminum, Delrin, and resin) (Fig. 3C, Fig. S3B). Dispersions were non-normally distributed. A Kruskal-Wallis test detected a significant effect of material on the dispersion of the deformation distribution (p = 2.28 × 10^−46^). Delrin was excluded from this analysis, as the least encountered material and the only material not evenly distributed within every session in which it appeared. Post-hoc pairwise Wilcoxon rank sum tests between the included material groups (Bonferroni correction, ɑ = 0.05) found significant differences in median dispersion between silicone and each of the less sticky materials (silicone and aluminum, p = 2.42 × 10^−29^; silicone and resin, p = 1.61 × 10^−40^), as well as between the two less sticky materials (p = 5.34 × 10^−3^). The silicone group had greater median dispersion than each of the other materials, with 70.61% and 73.83% of the silicone dispersion values exceeding those of the aluminum and resin distributions, respectively (Fig. 3C; Table 1).

**Table 1.**
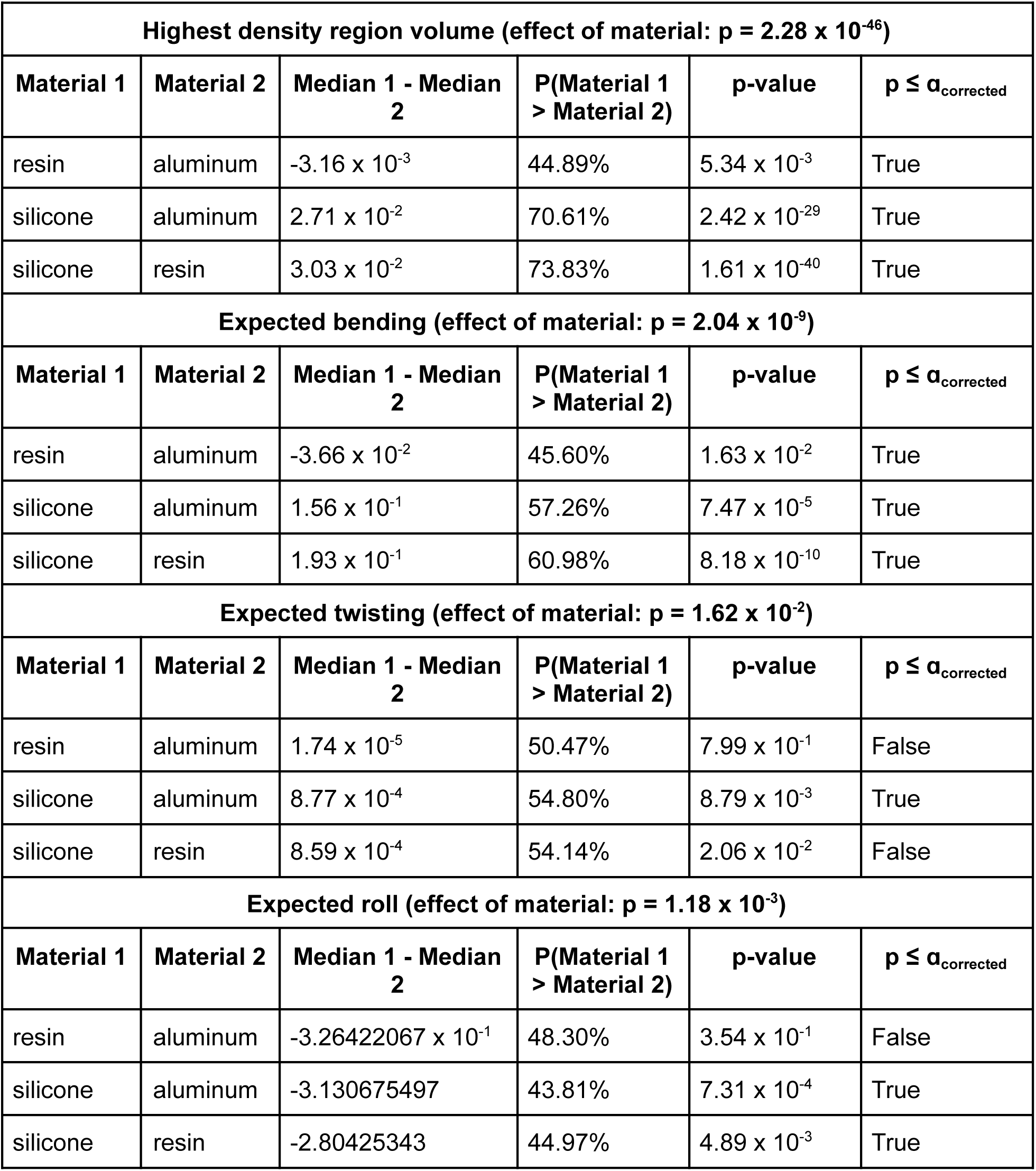
Comparison of the whisker deformation distribution’s HDR volume and mean between material groups.

Kruskal-Wallis tests on each dimension of the expected deformation (Fig. 3D) detected significant effects of material on all three (bending, p = 2.04 × 10^−9^; twisting, p = 1.62 × 10^−2^; roll, p = 1.18 × 10^−3^). Post-hoc pairwise Wilcoxon rank sum tests (Bonferroni correction, ɑ = 0.05) found significant differences in median expected bending and roll between silicone and each of the less sticky materials (bending, silicone and aluminum, p = 7.47 × 10^−5^; bending, silicone and resin, 8.18 × 10^−10^; roll, silicone and aluminum, p = 7.31 × 10^−4^; roll, silicone and resin, 4.89 × 10^−3^), a significant difference in median expected bending between resin and aluminum (p = 5.34 × 10^−3^), and a significant difference in median expected twisting between silicone and aluminum (p = 8.79 × 10^−3^). However, the probability of any dimension of the expected deformation for one material of a pair exceeding the other was generally lower than that observed in the comparisons of dispersion (Table 1). We conclude that the main effect of material variation on whisker deformation was an expansion of the whisker deformation distribution between the low stickiness (aluminum, resin) and high stickiness (silicone) surfaces, resulting in an increase in the sampling of high-strain whisker states.

### Whisking against the stickiest material evoked stronger but less frequent stick-slips

Stick-slip events are typical of frictional interactions between sliding surfaces. In whisker-based texture sensing, they have been observed as rapid transients in the angular or linear speed and acceleration of whiskers tracked using linear CCD arrays or single-view video (Arabzadeh et al., 2005; Isett et al., 2018; Ritt et al., 2008; Wolfe et al., 2008). These events are thought to be an important basis for whisker-based coarseness perception, with the rates and amplitudes of stick-slips observed to increase with grating spatial frequency (Isett et al., 2018) and sandpaper grain size (Wolfe et al., 2008) – both of which can be interpreted as increasing surface coarseness, as the whisker’s encounters with high-friction topographical features (grating edges and large grains) increase. Given these observations, we anticipated that stick-slips could also be affected by other aspects of the frictional interactions between whisker and surface, including surface stickiness.

Stick-slip events have not previously been observed in three dimensions. To identify and quantify these events analogously in 3D to previous lower dimensional descriptions, we used a directionless and whisker-centered set of variables to quantify whisker kinematics: the angular speed of a point on the whisker was the magnitude of the angular velocity of the vector from the whisker base to that point, and we derived the point’s angular acceleration and jerk from its angular speed. Stick-slips appeared as sharp whisker tip acceleration transients followed by sustained high-speed sliding of the whisker over the surface (Fig. 4A-F; Fig. S4; Supplemental Videos 5-8).

**Figure 4.**
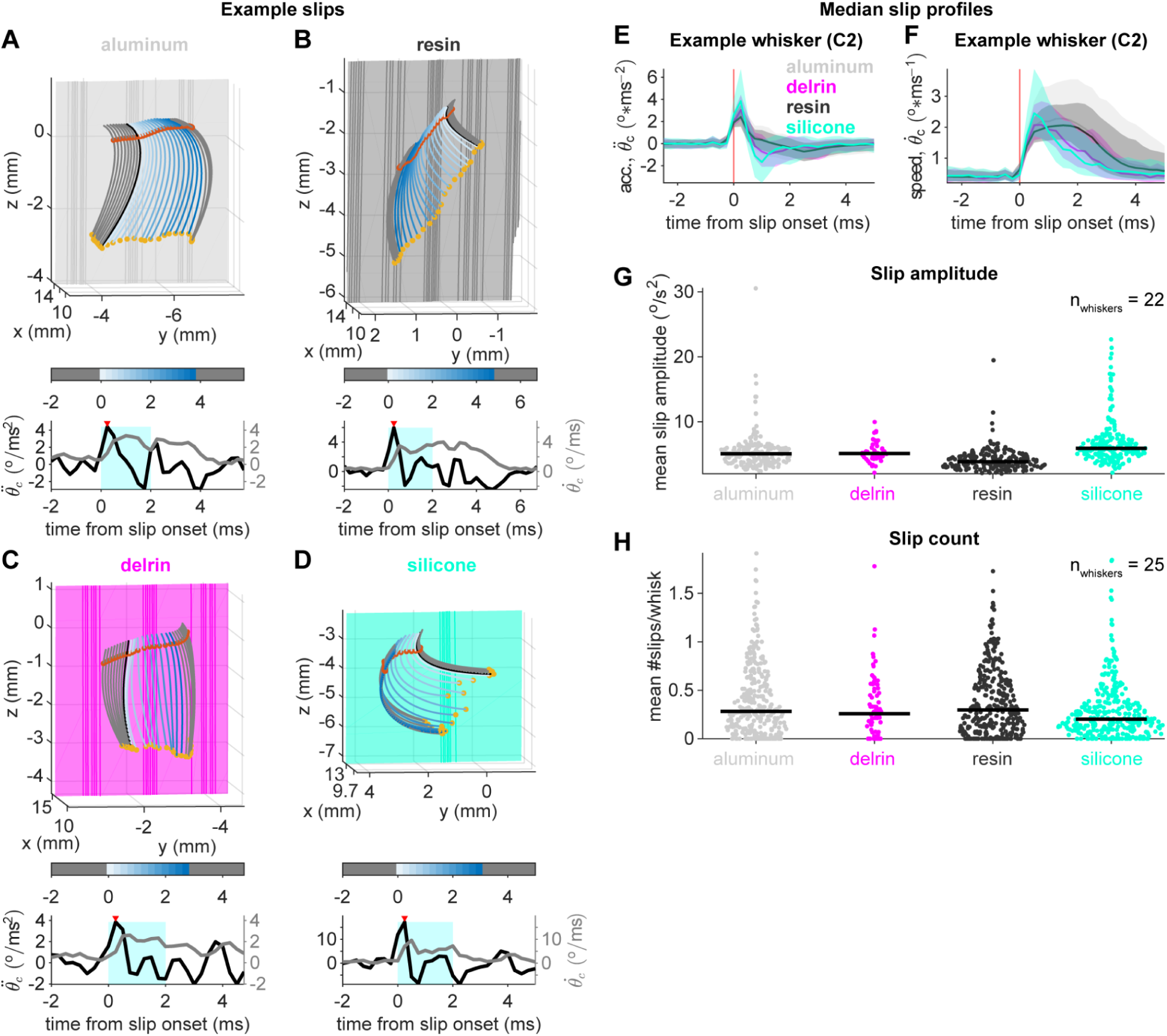
Whisking against silicone evoked higher-amplitude stick-slips than whisking against less sticky materials, but lowered stick-slip rates. **A-D)** Example slips against all materials presented. Upper panels: reconstructions of the whisker and surface at slip onset (black curve), over the duration of the slip (white to blue curves), and over the 2 ms preceding and following the slip (gray curves). Colored meshes are textured tile surfaces. View is zoomed in to the portion of the whisker from tip to reference point. Reference point (orange dot) and contact point (yellow dot) are shown for each frame. Lower panels: whisker tip acceleration (black trace) and speed (gray trace) profiles for the period from 2 ms before to 2 ms after the slip onset. Cyan shading indicates the region in which the slip peak acceleration is sought; red triangles mark the identified peak acceleration. **E, F)** Median acceleration (E) and speed (F) profiles of slips by the example C2 whisker in A-D, grouped by material. **G, H)** Scatter plots of mean slip amplitude (G) and rate (H), grouped by material. As in Figure 3C, each point corresponds to a single whisker instance observed under a single experimental condition.

Our analysis of stick-slip events focused on whiskers C2 and C3, as the whiskers best oriented to make large sweeps across the presented surfaces. As a kinematic description of stick-slip strength comparable to those used in previous studies (Isett et al., 2018; Wolfe et al., 2008), we defined the amplitude of a stick-slip as the peak tip acceleration reached within 2 ms of onset (Fig. 4A-D). We quantified stick-slip rate as the event count per whisk cycle. For each whisker instance, we computed the mean slip rate and amplitude in each experimental condition it encountered (Fig. 4G,H).

Kruskal-Wallis tests detected significant effects of material on stick-slip amplitude (p = 7.32 × 10^−34^) and rate (p = 1.40 × 10^−3^) (Table 2). As in the previous section, Delrin was excluded from the tested material set. Post-hoc pairwise Wilcoxon rank sum tests (Bonferroni correction, ɑ = 0.05) detected significant differences in median slip amplitude between all pairs of materials (resin and aluminum, p = 2.17 × 10^−12^; silicone and aluminum, p = 2.16 × 10^−9^; silicone and resin, p = 1.07 × 10^−32^). Median slip amplitude was greater in the silicone group than in the aluminum and resin groups, and greater in the aluminum group than in the resin group, with 67.29% and 83.75% of mean slip amplitude values in the silicone group exceeding those in the aluminum and resin groups, respectively, and 70.24% of mean slip amplitudes in the aluminum group exceeding those in the resin group (Table 2). Post-hoc pairwise Wilcoxon rank sum tests detected significant differences in median slip rate only between silicone and each of the less sticky materials (silicone and aluminum, p = 1.54 × 10^−3^; silicone and resin, p = 2.16 × 10^−3^), with lower median slip rate in the silicone group than in each of the other material groups, and with 57.38% and 56.98% of mean slip rate values in the aluminum and resin groups, respectively, would exceed those in the silicone group (Table 2). These results indicate that whisking against silicone evoked stronger but fewer stick-slip events than against the less sticky materials, suggesting that its greater stickiness may have allowed greater accumulation of strain in the elastic whisker prior to the release of slipping.

**Table 2.** Comparison of mean slip amplitude and rate between material groups.

| Mean slip amplitude (effect of material: $p = 7.32 \times 10^{-34}$) | | | | | |
| --- | --- | --- | --- | --- | --- |
| Material 1 | Material 2 | Median 1 - Median 2 | P(Material 1 > Material 2) | p-value | $p \leq \alpha_{\text{corrected}}$ |
| resin | aluminum | -1.23 | 29.76% | $2.17 \times 10^{-12}$ | True |
| silicone | aluminum | $8.30 \times 10^{-1}$ | 67.29% | $2.16 \times 10^{-9}$ | True |
| silicone | resin | 2.06 | 83.75% | $1.07 \times 10^{-32}$ | True |
| Mean slip rate (effect of material: $p = 1.40 \times 10^{-3}$) | | | | | |
| Material 1 | Material 2 | Median 1 - Median 2 | P(Material 1 > Material 2) | p-value | $p \leq \alpha_{\text{corrected}}$ |
| resin | aluminum | $1.45 \times 10^{-2}$ | 49.48% | $8.23 \times 10^{-1}$ | False |
| silicone | aluminum | $-8.18 \times 10^{-2}$ | 42.62% | $1.54 \times 10^{-3}$ | True |
| silicone | resin | $-9.63 \times 10^{-2}$ | 43.02% | $2.16 \times 10^{-3}$ | True |

In contrast, grating spatial frequency had limited effect on the rates and amplitudes of stick-slip events (Fig. S5A, B): Kruskal-Wallis tests detected a significant effect of spatial frequency only on slip amplitude (p = 9.48 × 10^−3^), with post-hoc pairwise Wilcoxon rank sum tests detecting no significant differences in slip amplitude between pairs of frequencies. This was unexpected, given past observations of the relationships between surface coarseness and stick-slip rate and amplitude (Isett et al., 2018; Ritt et al., 2008; Wolfe et al., 2008). We speculated that our observations could be a consequence of the uniformly low relief of the gratings we used, which was comparable to the grain diameter of a coarse sandpaper (Morita et al., 2011). To test this, we fabricated an additional set of silicone and resin gratings at the same spatial frequencies as previously used, with a new amplitude of 0.8 mm peak-to-trough. This amplitude exceeded the original value by a factor of 6.4, and was identical to that used by Isett et al. (Isett et al., 2018) for gratings of some of the same spatial frequencies. We presented these surfaces to the C1-C4 whiskers of two additional mice. Whisking against the higher-relief set of gratings evoked stick-slips of greater amplitude than the original lower-relief set (Fig. S5A, C). However, the effects of grating frequency on slip rate and amplitude remained weak: Kruskal-Wallis tests detected a significant effect of grating frequency on slip rate (p = 2.13 × 10^−2^) but not slip amplitude, and post-hoc pairwise Wilcoxon rank sum tests detected no significant differences in slip rate between pairs of grating frequencies. We conclude that while effects of grating frequency on stick-slip rate and amplitude are present, they may be overwhelmed when texture variation is multidimensional.

### Single-unit selectivity for the stickiest material increased with sensitivity to whisker strain

We recorded the activity of 10 TG units responsive to whisker touch during whisking against surfaces of varying texture. Whiskers in the C row were sought preferentially, as the C row was best oriented for tracking in both views and extensive contact with surfaces. However, one unit responsive to D1 whisker stimulation was also included. Based on their responses to manual whisker deflection, we identified 5 units as slowly adapting (SA) (Fig. 5A) and 4 as rapidly adapting (RA) (Fig. 5B). One unit’s response type was not identifiable (Fig. 5C). Several SA units showed marked increases in whisk-average spike rate during whisking against silicone compared to the less sticky materials (aluminum, Delrin, and resin). To quantify material discriminability, we computed the AUC for discrimination based on whisk-average spike rate between each pair of materials and between silicone and all other materials collectively. The spike rates of 5 of the 10 units significantly discriminated silicone from at least one other material (95% confidence interval of the AUC not including 0.5) (Fig. S6), and the spike rates of 4 of the 10 units significantly discriminated silicone from all other materials the unit encountered (Fig. 5D).

**Figure 5.**
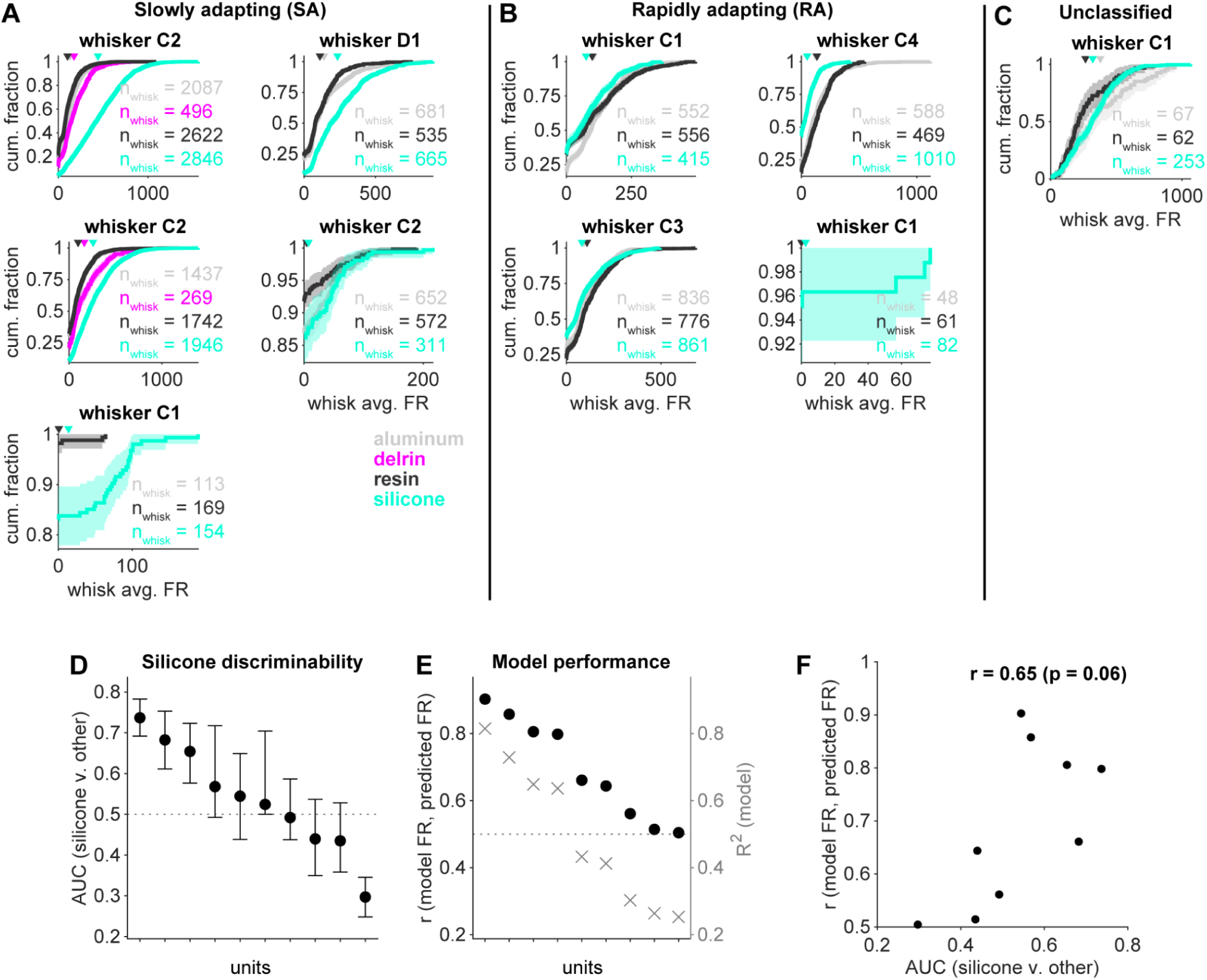
Some whisker follicle innervating units responded preferentially to whisking against silicone. **A - C**) Cumulative distributions of whisk-average spike rates for all units. Whisks included are restricted to those that contain whisking with contact. **D**) AUC for discrimination of whisks against silicone from whisks against other materials, based on whisk-average spike rate for each unit. Error bars indicate 95% CI. **E**) Performance for each unit of an ensemble of regression trees fitted to predict single-unit spike rate from smoothed bending, twisting, roll, and their first derivatives, quantified in terms of correlation between predicted and measured spike rate (r) and variance explained (R^2^). **F**) Model performance from panel E plotted against AUC for discrimination of whisks against silicone from panel D.

We took whisker bending, twisting, roll, and their first derivatives as the dimensions of the stimulus space to which each neuron was responding, based on previous observations of whisker bending and its rate of change as strong predictors of whisker follicle mechanosensory neurons’ activity (Bush, Schroeder, et al., 2016; Campagner et al., 2016; Severson et al., 2017). Regression analyses confirmed that the spike rates of the units we observed were generally explainable by this set of predictors (Fig. 5E). Given the expansion of the whisker strain distribution during whisking against silicone compared to the less sticky materials, we asked whether the discriminability of silicone from other materials based on a unit’s spike rate correlated with the sensitivity of that unit to whisker strain, quantified as the performance of the bending-, twisting-, and roll-based linear regression model in predicting its spike rate. We observed a positive correlation (r = 0.65, p = 0.06) between model performance and AUC for discrimination of silicone from all other materials (Fig. 5F). That the discriminability of silicone from less sticky materials based on neuronal response increased with units’ sensitivity to whisker strain, which reached higher magnitudes during whisking against silicone, suggests that strain in the whisker shaft is a viable physical basis for whisker-based stickiness perception.

## Discussion

We provide a first description of the whisker-surface interactions associated with whisking against surfaces of varying stickiness. To reach a general understanding of the representation of touch in the whisker system, characterization of whisker interactions with surfaces comprising a wide variety of material and topographic properties is needed.

Our results demonstrate that the sampling of high-strain whisker states and the strengths and rates of stick-slip events differed between whisking against high and low stickiness materials, providing multiple potential means of extracting surface stickiness information from whisker touch. These physical differences in the whiskers’ interactions with surfaces were reflected in selectivity for whisking against the high stickiness material in a subset of whisker follicle mechanosensory neurons. Taken together, these observations indicate that signals associated with the stickiness of surfaces whisked against are available to the whisker somatosensory system.

Although we have established a physical and a low-level neuronal basis for whisker-based stickiness sensing, many details of the associated perceptual experience remain unexplored. Our observations suggest that the material set used here may be discriminable into high stickiness (silicone) and low stickiness (aluminum, Delrin, and resin) groups. However, whether mice can report the relative stickinesses of surfaces based on whisker touch alone, whether and how stickiness interacts with coarseness and compliance during texture discrimination, and how the perceptual discriminability of surface stickiness relates to the coefficients of friction between whiskers and surfaces are questions for future work.

Although in this study we treat material identity as a categorical proxy for surface stickiness, stickiness as a continuous variable is likely best captured by the coefficients of friction between a sensor (such as a whisker or fingertip) and a contacted surface. The similarity of whisker–surface interactions and evoked neuronal responses among the low-stickiness materials we used in this study, which were intended to sample both low and intermediate stickiness, was unexpected; we speculate that these materials were too similar in their coefficients of friction with the whisker to meaningfully alter the strength of the frictional forces produced. Coefficients of friction are empirically determined, and while they have been measured for large libraries of pairs of materials with industrial applications, we are not aware of such measurements involving the whiskers or other hair of mice or rats. Determining coefficients of friction between whiskers and various materials is an important area for further investigation.

We have made the MATLAB-based 3D whisker tracking system we developed here available for broader use, as an extension of the 2D Whisker code library (Pammer et al., 2013). Like the original Whisker system, it takes the outputs of the Janelia whisker tracker (Clack et al., 2012) as its source of 2D whisker coordinates and initial identities; however, it can be adapted to read analogous information from output files produced by other 2D trackers, which may offer improved performance as new tools such as WhiskEras (Betting et al., 2020; Arvanitis et al., 2022) continue to be developed. The 3D tracking system is accompanied by a batch processing pipeline for analysis of multiple trials or sessions, which can be run locally or on a computing cluster; an annotation GUI for alignment of data to metadata; and a tile-fitting GUI for the semi-automated reconstruction of tile surfaces from silhouettes. Currently, the system can be used to track multiple whiskers in contact with a textured tile or in the absence of a defined stimulus object. We have applied it here to awake whisking against surfaces, awake whisking without contact, and passive whisker deflection by a handheld probe, and in other experiments to contactless whisker motion induced under anesthesia by optogenetic stimulation of the whisker pad (Kim et al., 2024).

This is not the first open-source 3D whisker tracking tool available to the field, and it differs from an existing tool, WhiskerMan (Petersen et al., 2020), in its strengths, limitations, and best use cases. In particular, we note that WhiskerMan’s manual error correction feature offers a significant advantage when manual curation of whisker fits is feasible, while the 3D extension of the Whisker library accommodates non-planar whisker shapes and reconstructs the full length of the whisker. We consider the 3D Whisker system best suited to applications involving high data volumes and complex whisker–surface interactions, such as stick-slips and contact-induced twisting, while WhiskerMan may be the more appropriate tool when the whisker does not twist and the state of its tip does not need to be considered.

## Code availability

Code for the reproduction of figures is available on Github (https://github.com/oconnorlab/Wyche_et_al_2026). The three-dimensional whisker tracking extension of the “Whisker” MATLAB code library and its associated data processing pipeline, including the annotation and surface fitting GUIs, are also available on Github (https://github.com/oconnorlab/Whisker3D).

## Data availability

Data for the reproduction of figures and example raw data for processing by the 3D Whisker tracking system are available through the Johns Hopkins Research Data Repository at https://archive.data.jhu.edu/previewurl.xhtml?token=58d85f0a-8b95-4464-ba8d-d2d9312ebeb0 (Wyche et al., 2026). All other data is available on request.

## Materials and Methods

All procedures were performed in accordance with protocols approved by the Johns Hopkins University Animal Care and Use Committee.

### Surgical procedures

Mice were implanted with titanium headcaps for head fixation during training and experiments, as described in previous studies (Severson et al., 2017). Between 4 hours and 1 day prior to the initiation of electrophysiological recordings, two craniotomies (2 mm medial-lateral by 0.5 mm anterior-posterior) were opened at 0 mm and 1 mm anterior and 1.5 mm lateral to bregma (Severson et al., 2017). Prior to and between experiments, craniotomies were protected with Body Double silicone elastomer and a thin layer of dental cement.

### Electrophysiology

Extracellular recordings in the whisker region of the left trigeminal ganglion were performed using 2 MΩ tungsten microelectrodes (World Precision Instruments or A-M Systems) as previously described (Severson et al., 2017). During recording, craniotomies were covered with saline and (for stabilization in some sessions, once a well-isolated unit was identified) 1% agarose in saline. The amplifier headstage was grounded to a silver chloride bead (World Precision Instruments) immersed in the saline bath or embedded in the agarose gel. Electrophysiological data was acquired using a World Precision Instruments DAM-80 amplifier at a sampling rate of 20kHz, with 10,000x gain and bandpass filtering between 300 and 3,000Hz.

### Spike sorting

Spike sorting was performed manually using MClust 4.4 (A.D. Redish, https://github.com/adredish/MClust-Spike-Sorting-Toolbox).

### Whisker and fur trimming

The fur on the left side of the face and neck, whisker rows A and E, the Greek whiskers, the microvibrissae, and whisker arcs rostral to 4 of rows B, C, and D were all trimmed close to the skin under isoflurane anesthesia. Whiskers and fur were initially trimmed up to 24 hours prior to the first recording session in each mouse, with further maintenance trimming as needed at the beginning (an hour or more before data acquisition) or end of experiment days. In experiments not including electrophysiology, the B and D rows were also fully trimmed before the first recording session; in experiments including electrophysiology, B and D row whiskers were retained as references in locating the C row region of the trigeminal ganglion, then trimmed immediately before initiating data acquisition.

### Water restriction

Mice were water-restricted to 1.5 mL per day for 3-7 days before training in head-fixed locomotion, and were maintained at no less than 80% of their initial weight for the duration of training and experiments. Mice were weighed before and after each training session or experiment, and supplementary water was given to ensure a minimum daily fluid consumption of 1.5 mL.

### Behavioral training

Mice were habituated to handling and then to head fixation prior to the initiation of treadmill training. Training began with small manual movements of the treadmill belt by the trainer, coaxing the mouse to run. A rotary encoder mounted on the rear axle of the treadmill detected movement. Once the mouse began to lick in expectation of reward during self- or trainer-initiated movements, the cumulative running time required to obtain a reward was gradually increased until reaching at least 2.5 seconds. Performance was considered sufficient for data acquisition when mice reliably self-initiated multi-second bouts of running over sessions of up to 45 minutes’ duration, with a rewarded running duration of at least 2.5 seconds. This was typically achieved within 1-2 weeks of training.

### Stimuli

Textured surfaces were vertical gratings of fixed amplitude (0.125 mm peak-to-trough for the low-relief set, and 0.8 mm peak-to-trough for the high-relief set), waveform shape (rectangular), and duty cycle (50%). Materials used were aluminum; Delrin acetal homopolymer (“Delrin”); Formlabs Black V4 methacrylate photopolymer (“resin”); and BBDINO Super Elastic platinum silicone (“silicone”). Aluminum and Delrin tiles were fabricated through CNC milling. Resin tiles were printed using a Formlabs 3B SLA printer from meshes generated in Blender 2.93, and silicone tiles were cast in Formlabs Black V4 resin molds that were also printed from meshes generated in Blender 2.93.

### Stimulus presentation

Each block of an experiment included three tiles having different combinations of material and grating frequency, which were mounted in a rotating three-slot stimulus presenter. Tiles could be presented in any of the four positions that were set for each session. Within a block, the identity of the tile presented and the position at which it was placed were randomly selected from trial to trial; each unique combination of tile identity and position was one “state”. States were removed from the set to be randomly sampled over when 5 total seconds of running in that state had accumulated. If the mouse was running at the end of a trial, the next trial would present the same tile in a random position if possible, to minimize the opportunity cost of the large movements required to change the presented tile. A block ended when all states were completed, and for the next block a new triplet of tiles was mounted on the stimulus presenter by the experimenter. Experiments ended when all blocks were completed, when the isolated unit was lost (in experiments including electrophysiology), or when the mouse ceased consuming rewards or initiating trials.

To minimize the acquisition of non-whisking frames, trials were triggered by the detection of briefly sustained treadmill movement. Exceptions to this condition on trial initiation were “blank” trials – in which no tile was presented – inserted into the sequence at 10- or 20-trial intervals, and manual stimulation trials in which the experimenter deflected the whiskers with a handheld probe. In these trials, data acquisition began immediately at the end of the 1 second inter-trial interval.

Data acquisition and stimulus presentation were centrally coordinated using a serial communication-based behavioral control system (Xu et al., 2022).

### Video acquisition

We used Streampix 9 software (Norpix) to record video over a 1,216 x 600 pixel ROI (square pixels, pixel size 10 μm) at a frame rate of 4,000 fps and an exposure time of 90 μs, using an Optronis Cyclone 2-2000 monochrome camera with a telecentric lens (Edmund Optics 0.25X 2/3” GoldTL Telecentric Lens) under 850 nm wavelength LED lighting. Frame capture was hardware-triggered, with frame triggers generated by Wavesurfer (https://wavesurfer.janelia.org). To achieve even background illumination throughout the ROI, the LED light path passed through an aspheric condenser lens followed by a stretched parafilm diffuser sheet. Prior to whisker tracking, videos were converted losslessly from raw Streampix SEQ format to TIF format, and were cropped during conversion to split the top-down and rear view portions of the ROI into separate files.

### Stereo calibration

Ground truth point correspondences between the top-down and rear views were obtained from images of a checkerboard calibration object, which was printed on a transparent acetate sheet and translated throughout the imaged volume on the same Zaber linear translation stages used to present the textured tiles. We used these points to estimate an affine fundamental matrix and affine camera matrices for the two views. To obtain reconstructions accurate to within a metric transform of ground truth, we took a stratified reconstruction approach based on the nominal dimensions of the checkerboard and the displacements in space between the set translation stage positions at which it was imaged. These two sets of metric constraints on the reconstruction allowed the estimation of a homography transforming the affine reconstruction to a metric one, and we computed metric camera matrices for the imaging system by applying this homography to the affine camera matrices. Reconstructions based on the metric camera matrices achieved sub-degree and sub-millimeter accuracy to the corner angles and grid step sizes of the true checkerboard, and corners of the reconstructed checkerboard reprojected to the original images with sub-pixel accuracy. This calibration process was implemented in MATLAB using algorithms detailed in *Multiple View Geometry In Computer Vision* (Hartley & Zisserman, 2003).

### 2D whisker tracking

Whiskers were tracked and initially classified in each view using the Janelia whisker tracker (Clack et al., 2012). Classification was subsequently corrected using code developed by Severson et al. (Severson et al., 2017), modified to accommodate this dataset.

### 3D whisker reconstruction

2D whisker coordinates outputted by the Janelia whisker tracker were corrected and cleaned before use in 3D whisker reconstruction: instances of whisker crossover in either view that caused the tips of adjacent whiskers to be swapped were automatically detected and corrected, and any portion of the tracked whisker that extended into the silhouette of the face or tile was excluded. Some crossovers could not be automatically detected for correction; trials heavily affected by these errors were later excluded based on manual review of trial-by-trial stick-slip event profiles, as uncorrected crossover errors were often detected as highly unnatural stick-slips.

Although the whisker offers few unambiguous points of correspondence between multiple views, point correspondences throughout the imaged space can be identified based on the epipolar geometry of the scene, which we estimated through the stereo calibration procedure. We identified corresponding points in the rear view for each point on each whisker in the top-down view using a custom stereo-matching procedure, and triangulated pairs of corresponding points into 3D space to reconstruct the tracked whisker centerline through a MATLAB implementation of the DLT algorithm for triangulation (Hartley & Zisserman, 2003). We then fitted degree 5 polynomial curves to the whisker centerlines. To standardize the orientation of the scene, curve fits were transformed by rotation and translation to a space with the whisker pad at its origin, oriented so that its x, y, and z axes (subsequently called the “world” coordinate system, {x_world_, y_world_, z_world_}) roughly aligned to the medial-lateral, rostral-caudal, and dorsal-ventral axes of the mouse (Fig. 2A-C).

### Whisker-centered coordinate frame

A static, whisker-pad-centered coordinate frame offers intuitively understandable representations of the whiskers’ and surfaces’ positions in space. However, the coordinate frame most relevant to the responses of follicle-innervating mechanosensory neurons is one that is fixed to the whisker follicle as it translates and rotates in the whisker-pad-centered space, as the neurons’ endings are (Huet et al., 2015; Bush, Solla, et al., 2016). This is the whisker-centered coordinate system. In keeping with the convention established by Huet et al. (Huet et al., 2015), we defined the whisker-centered coordinate system {x_0_, y_0_, z_0_} of each frame such that x_0_ lay antiparallel to the whisker base, y_0_ lay orthogonal to x_0_ in the proximal whisker shaft’s best fit plane of positive curvature (or, during contact, an estimate of this plane given the whisker’s position), and z_0_ lay orthogonal to both x_0_ and y_0_ (Fig. S7A, B).

### 3D surface reconstruction

We reconstructed tile surfaces from the edges of the tiles’ silhouettes in the two views. The tile mesh obtained for the first instance in a session of a given tile identity and position combination was re-used for all subsequent trials of matching tile identity and position.

User-specified initialization points, selected through a custom MATLAB GUI, guided the automated isolation of a minimum set of tile edges from among all edges detected in each image. The tile fitting protocol then automatically estimated the tile’s position and orientation in space based on the locations of the initialization points and the calibrated epipolar geometry of the scene. Slight buckling along tiles’ major axes could occur in the silicone and resin tiles; this was due to the elasticity of the cured silicone, and the plasticity of the resin in the period between initial printing and final thermal and UV curing. For tiles fabricated from these materials, deviations from planarity were fitted as quadratic functions of displacement along the tile’s horizontal and vertical edges. Finally, the terms of a transverse rectangular wave along the tile’s horizontal edge were fitted for all tiles. This semi-automated process produced tile surfaces with positions, orientations, buckle, and grating functions all fitted to the tile silhouettes.

#### Whisker shape computation

With the whisker’s centerline fitted as a continuously differentiable curve, its shape is fully described by its curvature (κ) and torsion (τ) as functions of arclength along the curve (Fig. S7C) (Kühnel, 2015). To summarize the overall shape of the whisker, we computed its total curvature and torsion by integrating over a fixed, reliably tracked span of its shaft, the reference ROI; the reference ROI extended from slightly distal to the whisker base to the arclength of the mid-shaft reference point. To measure the shape of the whisker in the top-down view alone, we computed the curvatures of reprojections of the 3D whisker curves into the top-down imaging plane, and integrated these values over the same ROI. As a complement to 2D curvature in the top-down view, we also computed the curvatures of reprojections of the 3D whiskers into a plane orthogonal to the top-down imaging plane, isolating the component of the whisker’s curvature that was not detectable in the top-down view.

### Whisker roll computation

Roll can be defined as change in the angle between the plane containing the whisker base – the best-fit plane to the proximal 60% of the whisker – and a chosen reference plane (Knutsen et al., 2008). What Knutsen et al. refer to as “torsion” or “torsional angle”, we describe here as “roll” to avoid confusion with torsion as a shape parameter. To measure the speed of the whisker’s roll out of the imaging plane, we used the top-down imaging plane as the reference plane roll was defined relative to; in all other analyses, the reference plane was the world x-y plane.

### Analysis of whisker strain

We defined whisker deformation, or the strain in the whisker shaft, as the contact-induced bending at the mid-shaft reference point p (Δκ_p_), the total contact-induced twisting (Δτ_total_) over the reference ROI, and the contact-induced axial roll of the whisker base (Δθ_roll_). Curvature deformation at the reference point provides a proxy for the transverse bending moments, transverse forces, and axial force applied to the whisker base during contact (Birdwell et al., 2007; Pammer et al., 2013); total torsion deformation over a portion of the shaft provides a proxy for the axial twisting moment about the whisker base.

Strain was defined relative to the whisker’s position-dependent intrinsic shape and orientation. We estimated intrinsic shape and orientation using polynomial functions of whisker base azimuthal angle (θ_az_), whisker base elevation angle (θ_el_), and (for intrinsic shape) arclength along the whisker (s), fitted to periods of low-acceleration whisking during blank trials with no surface presented (Fig. S7D, E, F).

We computed the strain probability distributions for each whisker instance from all frames meeting the inclusion criteria of whisker–surface contact (whisker tip within 0.8 mm of surface), active whisking (whisking amplitude at least 5°), and reliable whisker tracking (no outliers in whisker base angular speed, acceleration, or jerk, smoothed or unsmoothed whisker strain, or the first derivative of smoothed or unsmoothed whisker strain). Outlier speed was defined as exceeding the session 99.5^th^ percentile; outlier acceleration or jerk as falling outside the 0.25^th^ and 99.75^th^ percentiles; and outlier whisker strain or its first derivative as falling outside the 0.5^th^ and 99.5^th^ percentiles. Any bins of the distribution containing fewer than 10 samples were set to zero.

To obtain an ɑ% HDR volume for a distribution, we first computed the critical probability mass f_ɑ_ as the 100-ɑ percentile of the distribution’s probability mass values, then defined the ɑ% HDR as the set of all bins of the distribution having probability greater than f_ɑ_ (Hyndman, 1996). The volume of each HDR was then the number of bins in the HDR, normalized to the number of bins with probability greater than zero in any experimental condition.

Because the 50% HDR volumes and expected values of the strain distribution were non-normally distributed, we tested for effects of material and frequency using Kruskal-Wallis tests, with post-hoc pairwise rank sum tests if a significant effect of material was detected (Bonferroni correction, ɑ = 0.05). Delrin was excluded from the material set used in these tests. We computed the effect size for each comparison by normalizing the U statistic for the rank sum test to the product of the sample sizes of the groups compared.

### Coefficients of uncertainty

As a measure of the redundancy of a random variable X to a random variable Y, we computed the uncertainty of X given Y as the mutual information between X and Y normalized to the entropy of X.

### Stick-slip detection

We defined the onset of a stick-slip as a frame that was 1) an onset of whisker tip acceleration exceeding its high threshold (its 85^th^ percentile during whisking without contact), 2) preceded within 3 frames by tip jerk exceeding its high threshold (its 68^th^ percentile during whisking without contact), 3) preceded within 4 frames by a sustained period of tip speed below its low threshold (its 85^th^ percentile without whisking or contact), and 4) followed within 3 frames by sustained and overlapping periods of tip and reference point speed exceeding their respective high thresholds (85^th^ percentiles during whisking without contact) (Fig. S4). This set of criteria imposes the established definition of a stick-slip as a ballistic event consisting of an abrupt transition from a pinned to an unpinned, fast-sliding state of the whisker tip (Fig. 4E, F) (Ritt et al., 2008; Wolfe et al., 2008; Isett et al., 2018; Oladazimi et al., 2021), while dispensing with directional information that was not the focus of the present analyses.

### Analysis of stick-slip events

Analysis of stick-slips was limited to the C2 and C3 whiskers. We computed the mean slip rate and amplitude in each condition for each whisker instance, excluding conditions in which fewer than 10 slips or fewer than 10 whisks were observed. To test for effects of material and grating frequency on slip rate and amplitude, we performed Kruskal-Wallis tests with post-hoc pairwise rank-sum tests, again excluding Delrin from the material set considered.

### Firing rate regression analysis

We fitted ensembles of regression trees to predict single units’ smoothed spike rates (Gaussian kernel, standard deviation σ = 8 ms) from smoothed strain (Savitzky-Golay filter, degree 3, smoothing span 129 samples) and the first derivative of the smoothed strain, using MATLAB’s *fitrensemble* function.

## Acknowledgements

We thank Kyle Severson, Noah Cowan and Balazs Vagvolgyi for advice on analysis.

## Supplemental Information

**Figure S1.**
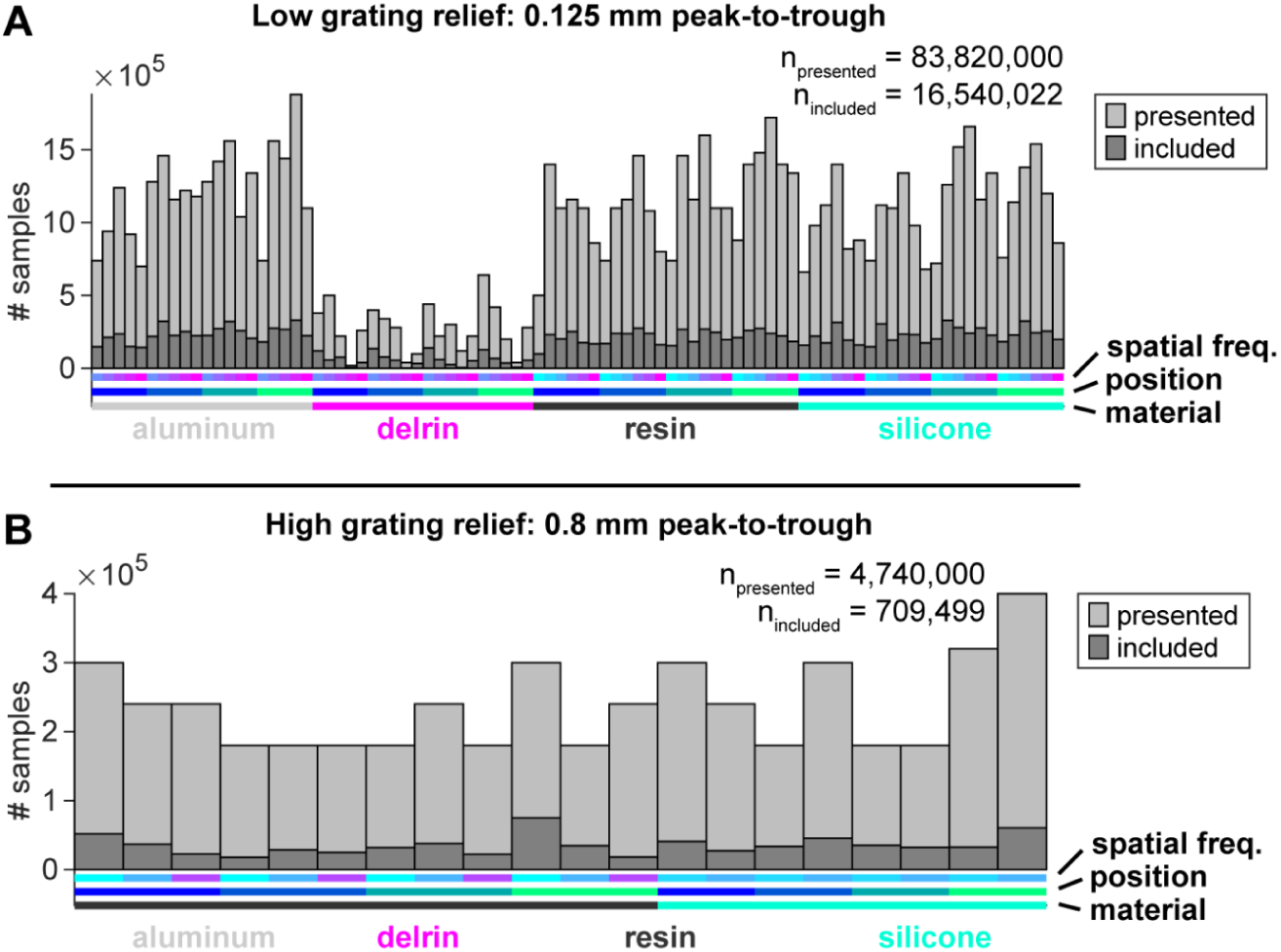
Samples were evenly distributed across experimental conditions, with the exception of Delrin as least-encountered material. **A, B)** Samples (frame-wise observations of single whisker instances) observed in each texture state, pooled across all sessions in which the original low-relief gratings (A) and the supplementary high-relief gratings (B) were used. “Included” samples are those passing the three inclusion criteria of surface contact, active whisking, and absence of outlier whisker strain or kinematics.

**Figure S2.**
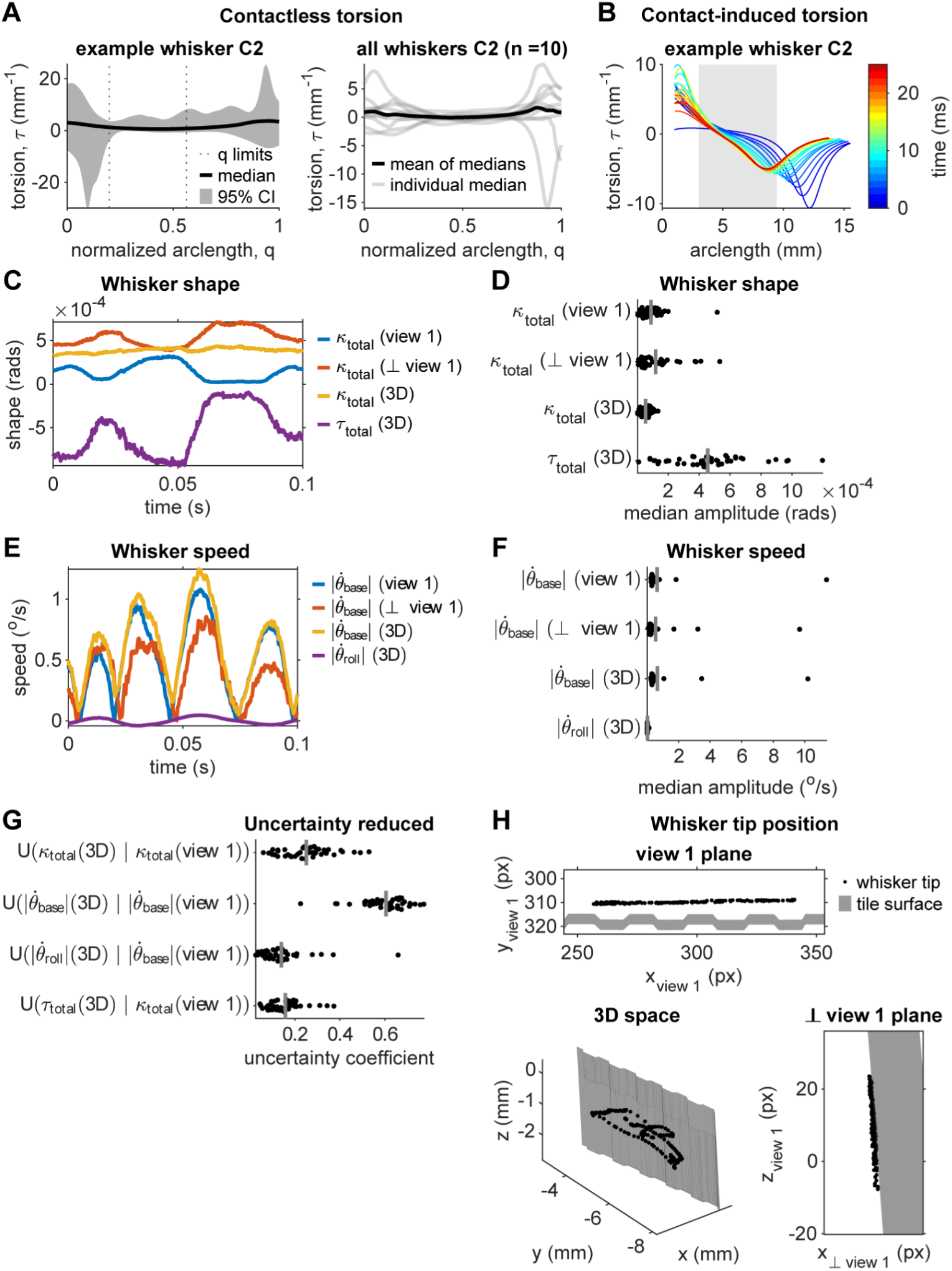
Information about whisker shape and motion during whisking against surfaces is lost when tracking is limited to a top-down view. **A)** Torsion in the shaft of a single example C2 whisker (left) and the full set of C2 whiskers (right), without contact or whisking. **B**) Torsion in the shaft of a single example C2 whisker over a 25 ms period of contact and whisking (with 5x temporal downsampling for visualization). **C**) Example 2D and 3D whisker shape signals measured during a 0.1 s period of contact and whisking. **D**) Median amplitudes of the whisker shape signals for all whisker instances. **E**) Example 2D and 3D whisker motion signals measured during the same period of contact and whisking as in C. **F**) Median amplitudes of the whisker motion signals for all whisker instances. **G**) Uncertainty coefficients between 3D whisker shape and motion signals and their 2D comparators. **H**) Position of the whisker tip over the same time period as in C and E, at 2x temporal downsampling.

**Figure S3.**
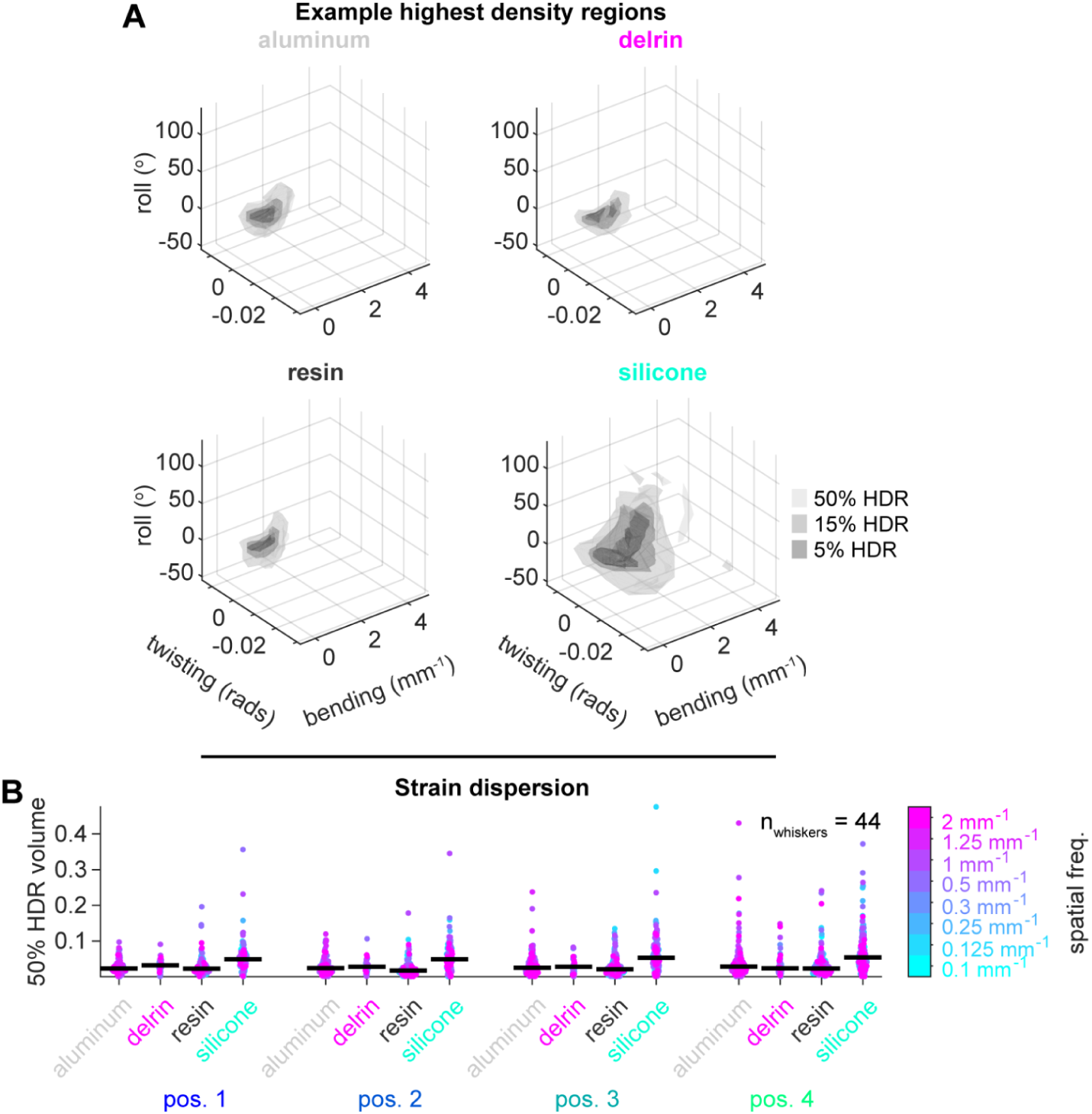
Highest density regions of the whisker deformation distribution. **A)** Gray volumes indicate the boundaries of the 50%, 15%, and 5% highest density regions of the example distributions shown in Figure 3B. **B)** Scatter plots of the 50% highest density region volumes of all whisker instances’ deformation distributions; points are grouped by position, subgrouped by material, and color coded by frequency. Each point corresponds to a single whisker instance that was observed under the conditions indicated.

**Figure S4.**
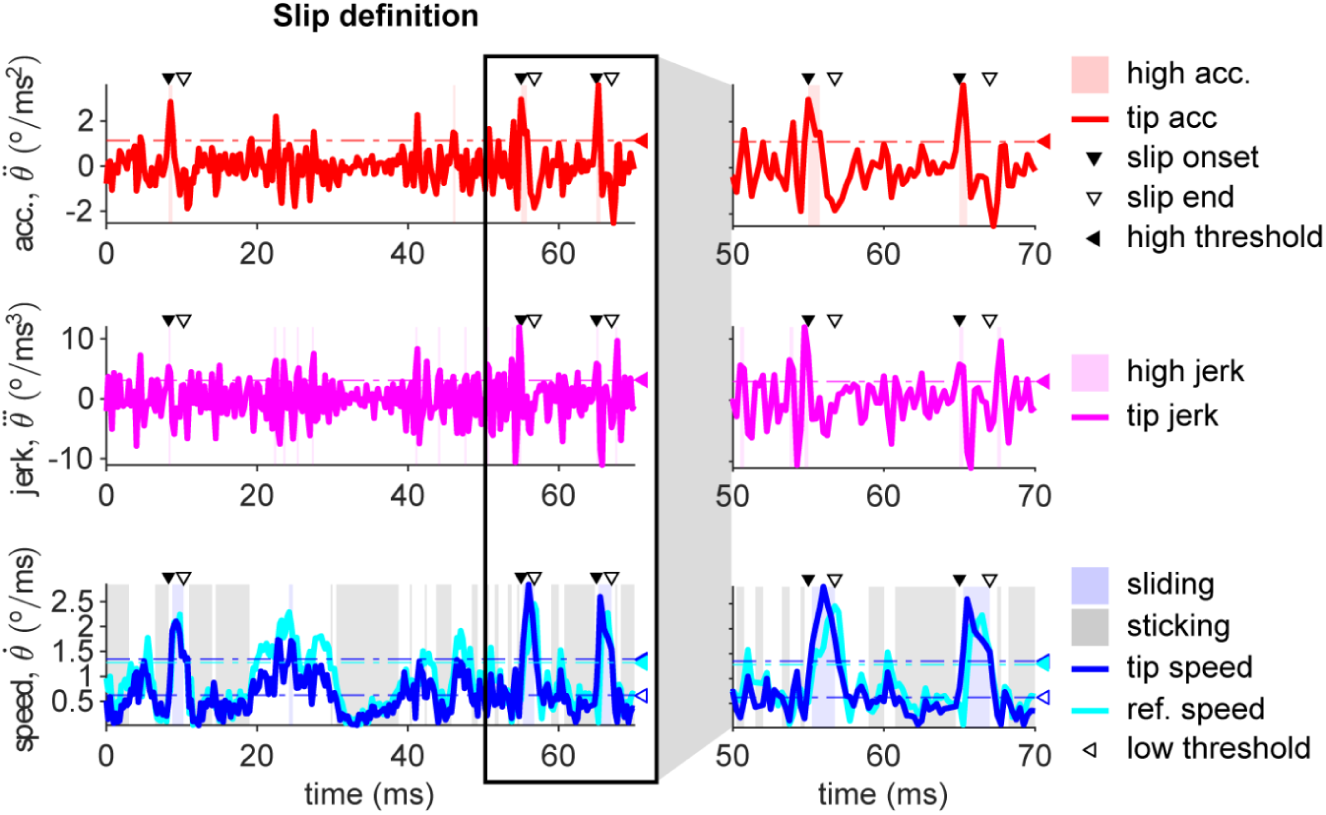
Stick-slip event boundaries were extracted from the kinematics of the whisker tip and reference point. Whisker tip acceleration (top), whisker tip jerk (middle), and whisker tip and reference point speed (bottom) traces over a random interval containing multiple slips. Filled and unfilled triangles above each panel indicate the start and end times, respectively, of the detected stick-slip events. Filled (all panels) and unfilled (bottom only) triangles to the right of each panel mark the high and low thresholds, respectively, for the plotted variables. Shading indicates frames classed as high acceleration, high jerk, sliding, or sticking, based on threshold crossings. Inset at right zooms in on two slips.

**Figure S5.**
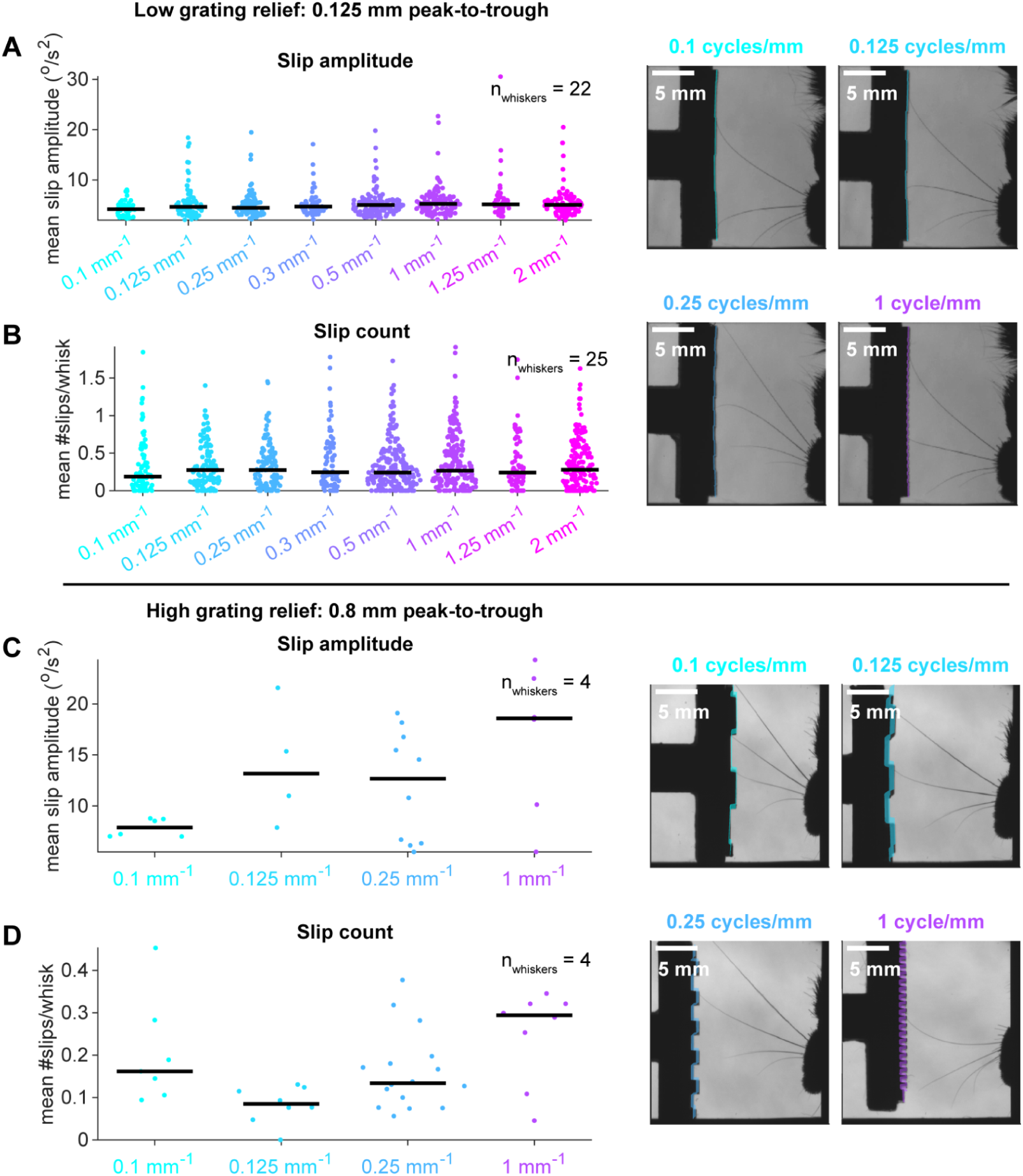
Relationships of stick-slip amplitude and rate to spatial frequency were weak in both high and low grating relief conditions. **A, B)** Scatter plots of mean slip amplitude (A) and rate (B), grouped by frequency, for the low-relief set of gratings. **C, D)** Scatter plots of mean slip amplitude (C) and rate (D), grouped by frequency, for the high-relief set of gratings. As in Figures 3 and 4, each point corresponds to a single whisker instance observed under a single experimental condition.

**Figure S6.**
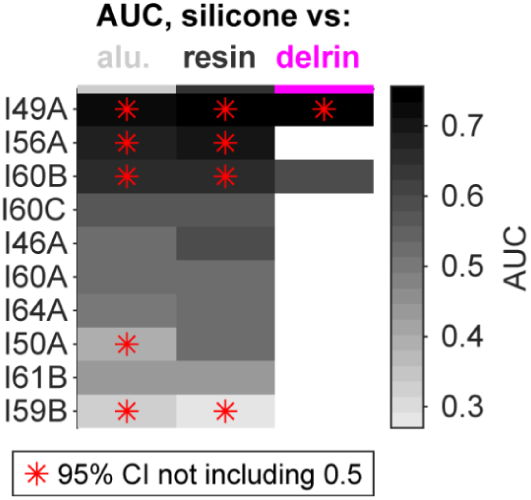
Whisks against silicone were discriminable from other materials on the basis of single-unit response. Heatmap displays, for each unit, the AUC for discrimination of silicone from each other material the unit encountered.

**Figure S7.**
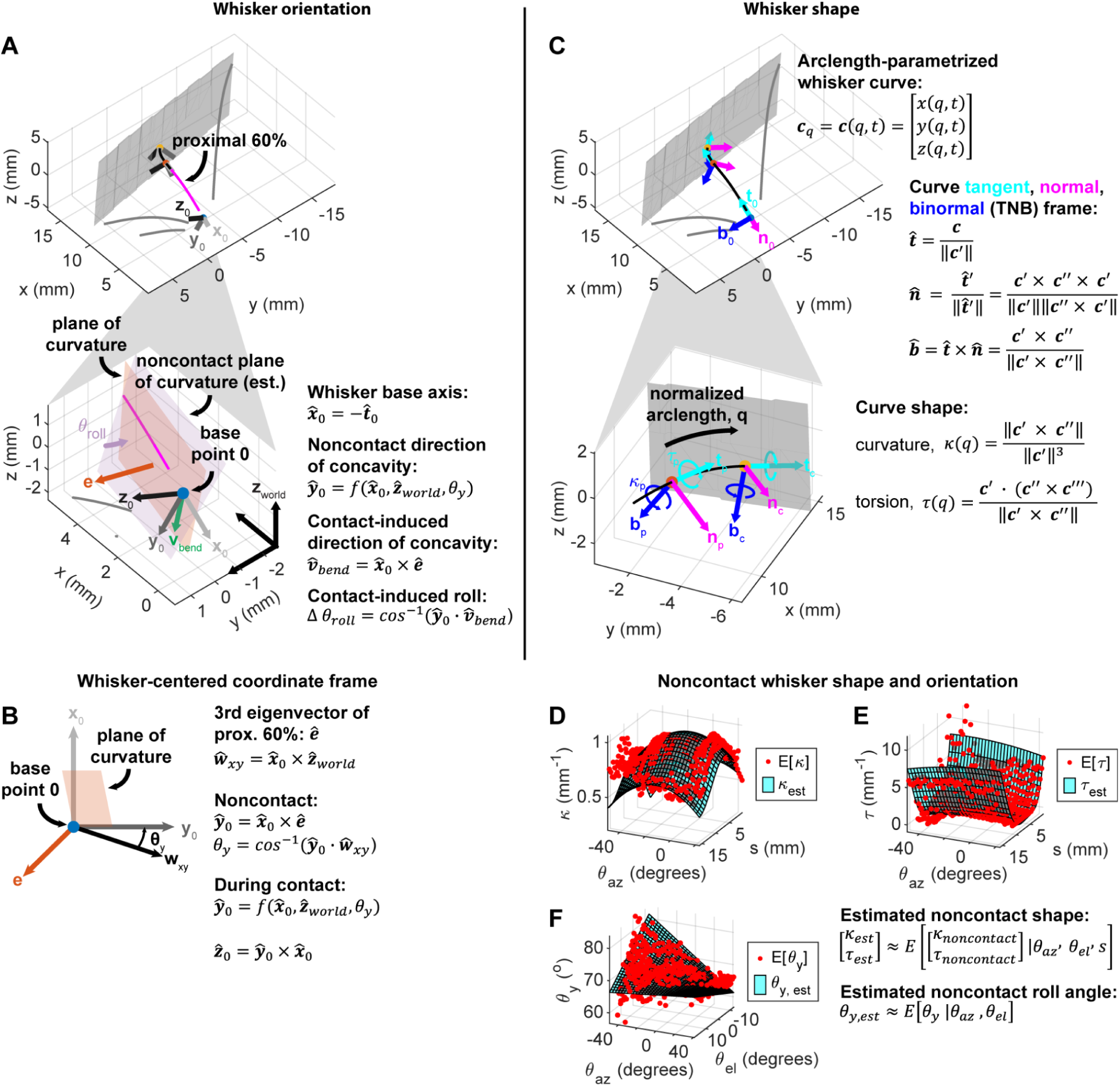
Computation of whisker shape, strain, and orientation. **A)** Contact-induced whisker roll was defined as the angle between the instantaneous plane of curvature of the proximal 60% of the whisker shaft (orange), and the estimated noncontact plane of curvature of the proximal 60% of the whisker shaft (purple). **B)** The whisker’s estimated noncontact plane of curvature defined the directions of the y and z basis vectors of the whisker-centered coordinate frame. **C)** The whisker curve’s tangent, normal, binormal (TNB) frame at each point along its length was computed from the curve and its first three derivatives. The curve’s shape (curvature and torsion) was the angular velocity of the TNB frame with respect to normalized arclength. **D-E)** The curvature (D) and torsion (E) of the undeflected whisker, given arclength along the whisker and the position (azimuth and elevation angles) of the whisker base, were estimated as fifth-order polynomial surfaces (cyan meshes) fitted to the average values observed during low-acceleration whisking without contact (red dots). Elevation angle is limited to a single value for visualization. **F)** The angle between the undeflected proximal whisker segment’s plane of curvature and the world x-y plane, given the whisker’s position, was estimated as a quadratic surface (cyan mesh) fitted to the average values observed during low-acceleration whisking without contact (red dots).

**Supplemental Video 1 – Example trial with aluminum tile and spikes from a whisker C2 slowly adapting unit.**

Video slowed 10x from data acquisition rate of 4,000 fps – 10x temporal downsampling, playback speed 40 fps. Whisker C2 shown in black, all other whiskers (C1, C3, and C4) in gray. For each whisker, blue marker indicates the whisker base; orange, the fixed-arclength reference point; and yellow, the whisker tip. Left column shows reprojections of the 3D scene reconstruction into the original top-down (upper) and rear (lower) views; right panel shows 3D scene reconstruction.

**Supplemental Video 2 – Example trial with Delrin tile and spikes from a whisker C2 slowly adapting unit.**

Same unit as in Supplemental Video 1. Video slowed 10x.

**Supplemental Video 3 – Example trial with resin tile and spikes from a whisker C2 slowly adapting unit.**

Same unit as in Supplemental Videos 1-2. Video slowed 10x.

**Supplemental Video 4 – Example trial with silicone tile and spikes from a whisker C2 slowly adapting unit.**

Same unit as in Supplemental Videos 1-3. Video slowed 10x.

**Supplemental Video 5 – Example stick-slip against aluminum tile.**

Same trial as in Supplemental Video 1 and same stick-slip as in Figure 4A. Video slowed 400x – no downsampling, playback speed 10 fps. All views (top-down, rear, and 3D) are zoomed in to the tip of the C2 whisker. The C2 whisker at the present time point is shown in black in each frame; as in Figure 4A-D, the C2 whisker at previous times is shown in gray (over the 2 ms preceding and following the slip) or in a gradient from white to blue (over the duration of the stick-slip).

**Supplemental Video 6 – Example stick-slip against Delrin tile.**

Same trial as in Supplemental Video 2 and same stick-slip as in Figure 4B. Video slowed 400x.

**Supplemental Video 7 – Example stick-slip against resin tile.**

Same stick-slip as in Figure 4C. Video slowed 400x.

**Supplemental Video 8 – Example stick-slip against silicone tile.**

Same trial as in Supplemental Video 4 and same stick-slip as in Figure 4D. Video slowed 400x.

